# Effects of 3-Nitrooxypropanol on Methane Emission, Growth and Feed Intake in Growing Calves from 5 Months of Age

**DOI:** 10.1101/2025.05.22.655456

**Authors:** Eline E. A. Burgers, André Bannink, Nicola Walker, Reto Zihlmann, Sanne van Gastelen

## Abstract

This study aimed to test the effect of 3-nitrooxypropanol (**3-NOP**) on methane (**CH_4_**) emission, feed intake, and body weight (**BW**) of calves from 5 mo of age. Seventy calves were enrolled in the acclimatization period (3 wk), after which 60 calves were selected based on their ability to use the automated feed bins (**FB**) and the GreenFeed system. These 60 calves were blocked in pairs for sex, breed, and BW, and within block randomly assigned to 1 of 2 dietary treatments: a partially mixed ration (**PMR**) supplemented with (1) a placebo contrate (**CTL**), or with (2) a 3-NOP concentrate (on average 121 mg 3-NOP/kg dry matter of total ration). The acclimatization period was followed by a covariate period (2 wk) in which baseline measurements were performed, and afterwards the treatment period (12 wk) started where the calves received 1 of the 2 dietary treatments. Next to PMR, the calves received up to 1280 g/ day bait in the GreenFeed system, which was used for emission measurements. For analyses, all data was averaged for the covariate period and for different periods in the treatment wk: 1-3, 4-6, 7-9 and 10-12. Upon 3-NOP supplementation, CH_4_ production (g/d) was significantly reduced by 31.3% in wk 1 – 3, by 27.1% in wk 4 – 6, by 30.0% in wk 7 – 9, and by 30.7% in wk 10 – 12 compared to CTL, and CH_4_ intensity (g/kg BW) was significantly reduced by 30.0% in week 1 – 3, by 25.8% in wk 4 – 6, by 27.9% in wk 7 – 9, and by 28.6% in wk 10 – 12 compared to CTL. Overall, 3-NOP calves had a reduction of 29.8% for CH_4_ production, 19.4% for CH_4_ yield (g/kg dry matter intake; **DMI**), and 27.9% for CH_4_ intensity. Calves receiving 3-NOP had a lower total DMI (6.1 kg/d) compared to CTL (6.9 kg/d), driven by a lower DMI of PMR. In every period, calves in 3-NOP had a lower BW (overall: 251 kg) compared to CTL (overall: 258 kg). The 3-NOP group and the CTL group did not differ in initial BW, but calves in the 3-NOP group had a lower final BW (290 kg) compared to CTL (301 kg). Both feed to gain ratio and feed efficiency did not differ between the two treatment groups. It can be concluded that 3-NOP is a promising strategy to persistently decrease CH_4_ emission in growing beef calves from 5 to 8 mo of age, without negatively impacting feed efficiency.

## Introduction

Beef cattle produce enteric methane (**CH_4_**) as a natural by-product of microbial fermentation in the rumen. This process allows them to effectively turn human inedible biomass, such as grass, into high quality protein in the form of meat for human consumption (Gerber et al., 2015). Enteric CH_4_ emission contributes to greenhouse gas (**GHG**) emissions, and estimates suggest that non-lactating and non-dairy cattle are the largest animal source of enteric CH_4_ emission in Europe followed by dairy cattle (FAOSTAT, 2019). Hence, enteric CH_4_ emission has become one of the main targets of GHG mitigation practices for the beef industry. Several dietary mitigation strategies exist to reduce enteric CH_4_ emission (Hristov et al., 2013). One of these strategies is a feed additive, 3-nitrooxypropanol (**3-NOP**; marketed as Bovaer®, dsm-firmenich AG, Kaiseraugst, Switzerland). Arndt et al. (2023) concluded that 3-NOP was the most effective CH_4_-mitigating feed additive within the category of rumen manipulation caused by feeding methanogen inhibitors. According to meta-analyses, 3-NOP effectively reduced enteric CH_4_ emissions in beef cattle by on average 22% to 35% (Dijkstra et al., 2018; Orzuna-Orzuna et al., 2024), which is a lower average reduction than reported for dairy cattle (−39%; Dijkstra et al., 2018).

In dairy cows, the efficacy of 3-NOP has been tested extensively, in many studies representing different stages of the lactation (e.g. Melgar et al., 2021; Schilde et al., 2021; van Gastelen et al., 2024). Also, several studies investigated the efficacy of 3-NOP for beef cattle in feedlots (e.g. Kim et al., 2019; Alemu et al., 2021; Almeida et al., 2023). Animals usually arrive at feedlots shortly after weaning (7 to 9 mo of age), as yearlings (12 to 18 mo of age) or at two and a half years of age. In the meta-analysis by Orzuna-Orzuna et al. (2024), the average body weight (**BW**) of the animals in the 15 included studies was 420 kg, indicating an age of about 12 mo.

Calves are born as non-ruminant animals, and from 2 days of age, microorganisms start colonizing the rumen of the calves (Rey et al., 2014; Li et al., 2023). Between 15 and 83 days of age, solid food intake increases rapidly, including roughage and concentrates, and enteric CH_4_ production will follow. It may thus be assumed that growing calves at 5 mo of age have a similar response to 3-NOP as older beef cows as evaluated by Orzuna-Orzuna et al. (2024). However, for young growing calves (i.e., <6 mo of age), studies of 3-NOP as a dietary additive to mitigate enteric CH_4_ emission are limited. When 3-NOP was supplied in early life, i.e. from birth until 3 wk post weaning, calves showed a persistent reduction in CH_4_ up to 12 mo of age (Meale et al., 2021). Only Kirwan et al. (2024) evaluated the effect of 3-NOP on enteric CH_4_ emission and performance in growing calves of less than 6 mo of age and concluded that CH_4_ emission was reduced by approximately 30%. Supplementation of 3-NOP did not affect dry matter intake (**DMI**), BW, or average daily gain (**ADG**). However, some other studies with growing beef cattle did report a reduction in DMI upon 3-NOP supplementation (Romero-Perez et al., 2014; Vyas et al., 2016), but no effect on performance. As such, the response of growing calves of less than 6 mo of age to 3-NOP supplementation in terms of CH_4_ emission, feed intake, and growth has not been completely elucidated yet.

Therefore, the aim of the current study was to test the efficacy and safety of 3-NOP for calves from 5 until 8 mo of age, and evaluate the effect on CH_4_ emission, feed intake, and BW. We hypothesized that 3-NOP supplementation would persistently decrease CH_4_ emission but would not affect feed intake and – despite the potential additional available energy due to the reduction of CH_4_ emission – also not positively affect growth performance.

## Materials and Methods

### Study Design

The study was conducted from June to October 2022 at the research facility of Wageningen Livestock Research (Dairy Campus, Leeuwarden, the Netherlands), under the Dutch law on Animal Experiments in accordance with European Union Directive 2010/63 and approved by the Animal Welfare Body of Wageningen Research (Lelystad, Netherlands). The study followed a complete randomized block design with 2 dietary treatments and 60 calves that were on average 5 mo of age at the start of the treatment period (5.1 ± 0.11 mo of age; mean ± SD). Each dietary treatment group consisted of 9 females (4 Belgian Blue and 5 Holstein Friesian) and 21 males (all Holstein Friesian). Part of the calves (n = 17, of which n=7 for the dietary control group and n=10 for the dietary treatment group) were born and housed on Dairy Campus until the start of the study, whereas the other calves (n = 43, of which n=23 for the dietary control group and n=20 for the dietary treatment group) were purchased from several Dutch farms and housed on a specialized calf rearing quarantine facility in accordance with the Dairy Campus procedures from 2 wk of age to 4 mo of age. This procedure was followed to ensure a minimum age difference between all calves (i.e., 2 wk), which would not have been possible when enrolling calves from Dairy Campus alone, whilst ensuring the strict health status requirements of Dairy Campus.

The study consisted of an acclimatization period, a covariate period, and a treatment period. In total, 70 calves were enrolled in the acclimatization period, which lasted for 3 wk. During the acclimatization period the calves could get used to the new barn and learn to use the GreenFeed system (C-Lock Inc., Rapid City, SD) and the automated Insentec feed bins (**FB**; Roughage Intake Control (RIC) system, Hokofarm Group BV, Marknesse, Netherlands). At the end of the acclimatization period, 60 calves were selected based on their ability to use the FB and GreenFeed system. These 60 calves were blocked in pairs for sex, breed, and BW, and within block randomly assigned to 1 of the 2 dietary treatments. The acclimatization period was followed by a covariate period of 2 wk in which the baseline measurements were taken. After the covariate period, the treatment period started, consisting of 12 wk during which the calves received 1 of the 2 dietary treatments.

During the entire study, calves were loose housed together in one pen consisting of a slatted floor at the feed aisle and the remainder being a deep litter area. The deep litter area was filled with fresh straw once weekly. The calves had access to 2 GreenFeed units and 12 FB and had free access to a water trough.

### Feeding and Dietary Treatments

During the covariate period, all calves received the same basal diet, fed as a partially mixed ration (**PMR**). After the covariate period, the calves were assigned to 1 of the 2 dietary treatments: where the PMR was (1) supplemented with a placebo concentrate (i.e., silicon dioxide + 1,2 propanediol; control [**CTL**]) or (2) with a 3-NOP concentrate (i.e., 10% 3-NOP on silicon dioxide + 1,2-propanediol). Both the CTL and the 3-NOP concentrate was added to the PMR to reach a target concentration of 175 mg CTL or 3-NOP/kg dry matter (**DM**) in the PMR, being 150 mg CTL or 3-NOP/ kg DM on complete ration level (including the non-supplemented GreenFeed bait). The use of 3-NOP in animal feed was approved by the Veterinary Drugs Directorate Division (Utrecht, Netherlands) prior to the start date. The chemical composition of the individual ration ingredients is presented in Table 1. The ingredient and chemical composition of the basal diet, CTL diet, and 3-NOP diet is presented in Table 2.

**Table 1.** Chemical composition (in g/kg DM, unless stated otherwise) of the individual feed ingredients.

|  | Maize silage | Lucerne | Concentrate |  |  | Soybean meal | GreenFeed bait<sup>4</sup> |
| --- | --- | --- | --- | --- | --- | --- | --- |
|  |  |  | Basal diet<sup>1</sup> | CTL<sup>2</sup> | 3-NOP<sup>3</sup> |  |  |
| DM (g/kg) | 355 | 914 | 889 | 884 | 884 | 877 | 883 |
| OM | 957 | 899 | 926 | 926 | 921 | 935 | 936 |
| CP | 65 | 172 | 149 | 135 | 143 | 464 | 122 |
| Crude Fat | 28 | 24 | 41 | 34 | 33 | 25 | 31 |
| Gross energy (MJ/kg DM) | 18.6 | 16.9 | 16.3 | 16.4 | 16.3 | 17.3 | 15.6 |
| NDF | 354 | 387 | 283 | 280 | 276 | 125 | 337 |
| ADF | 207 | 291 | 159 | 160 | 156 | 63 | 204 |
| ADL | 9.8 | 62 | 44 | 43 | 42 | 1 | 14 |
| Starch | 380 | 16 | 278 | 268 | 263 | 21 | 116 |
| Sugar | 0 | 51 | 33 | 31 | 30 | 100 | 92 |

<sup>1</sup> Ingredient composition (g/kg DM): palm kernel flakes = 297, maize = 284, wheat semolina = 120, wheat = 101, rumen protected rapeseed meal (Mervobest, NuScience) = 75, lupine = 41, CaCO<sub>3</sub> = 24, lucerne = 20, citrocol = 20, NaCl = 10, MgO = 5, and mineral premix = 2.
<sup>2</sup> Ingredient composition (g/kg DM): palm kernel flakes = 282, maize = 270, wheat semolina = 113, wheat = 104, rumen protected rapeseed meal (Mervobest, NuScience) = 71, lupine = 47, lucerne = 19, sugar beet pulp = 19, NaCl = 9, mineral premix = 8, and placebo = 58.
<sup>3</sup> Ingredient composition (g/kg DM): palm kernel flakes = 282, maize = 270, wheat semolina = 113, wheat = 104, rumen protected rapeseed meal (Mervobest, NuScience) = 71, lupine = 47, lucerne = 19, sugar beet pulp = 19, NaCl = 9, mineral premix = 8, and 3-NOP = 58.
<sup>4</sup> Ingredient composition (g/kg DM): sugar beet pulp = 300, soybean meal = 221, corn gluten meal = 87, wheat = 72, alfalfa = 70, sunflower kernel meal = 70, molasses = 50, barley = 50, corn = 49, rapeseed meal = 21, NaCl = 10.

**Table 2.**
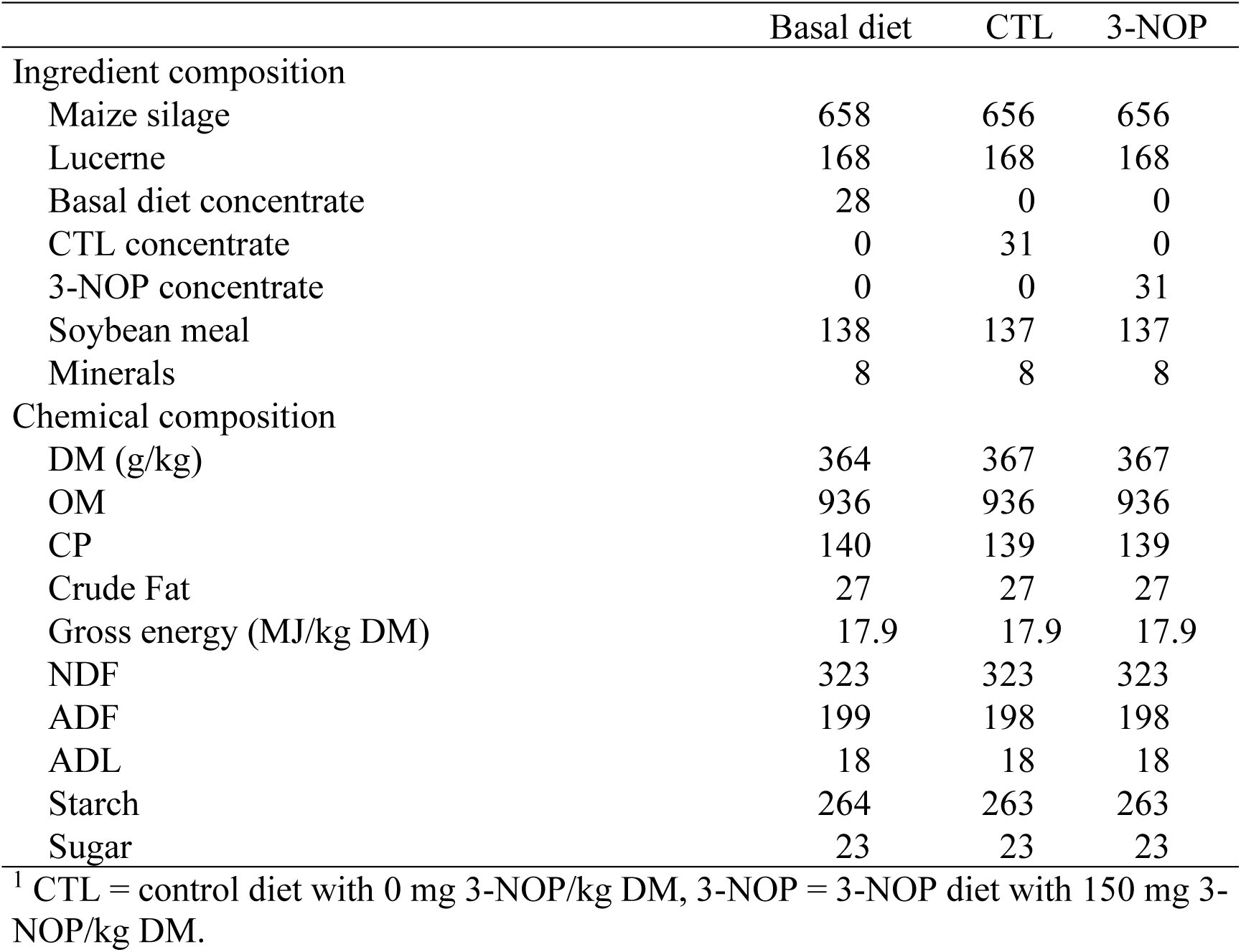
Ingredient and chemical composition (in g/kg DM, unless stated otherwise) of the diets, where GreenFeed bait is not included in the calculation^1^.

Feeding rate of all diets was adjusted daily to yield refusals equal to a target of 10% of intake. Fresh PMR was prepared once a day, by using a Trioliet feed mixing robot (Trioliet feeding technology, Oldenzaal, Netherlands). Prior to offering new feed, feed refusals were removed from the FB. Calves had access to all FB with the appropriate treatment diet, resulting in 60 calves having access to 12 FB during the covariate period or each group of 30 calves having access to their allocated 6 FB during the treatment period, always representing 5 calves per FB. Next to the PMR, the calves had access to 2 GreenFeed units, where they received a maximum of 1280 g of GreenFeed bait daily (chemical and ingredient composition shown in Table 1). This amount of GreenFeed bait was based on 4 GreenFeed visits per day, each with 8 cup drops of approximately 40 g bait per cup drop. As such, GreenFeed bait made up approximately 13% of the total DMI in both groups.

### Measurements

Individual DMI of the PMR was automatically recorded at each visit of the animal to the FB. The amount of GreenFeed bait received by each animal was also recorded in terms of the amount of cup drops of feed that the animal received at each visit. On a weekly basis throughout the study, the amount of bait delivered in each cup drop was weighed and tested for both GreenFeed units, to ensure accuracy on the calculation of DMI. Total DMI was calculated by adding the DMI of the PMR to the DMI of the GreenFeed bait on a daily basis.

The individual BW of each animal was manually recorded weekly by placing them on a weighing scale within the barn (Welvaarts Weegsystemen BV, ‘s Hertogenbosch, Netherlands). The following variables from these recordings were used in data analyses: weekly BW, initial BW (single measure at 1 week after start of the acclimatization period), final BW (single measure at the end of the treatment period), BW gain (final BW – initial BW), and ADG (BW gain / days[final – initial]). With the ADG, the feed to gain ratio was calculated by dividing average daily total DMI (kg) over ADG (kg), and the feed efficiency was calculated by dividing ADG (kg) over average daily total DMI (kg).

Gas emissions [CH_4_, hydrogen (**H_2_**), and carbon dioxide (**CO_2_**)] were measured on individual animal level by using the GreenFeed system as described by van Gastelen et al. (2022). All calves had free access to 2 GreenFeed units that were placed in the pen at the beginning of the acclimatization period and left there for the entire study. Gas (CH_4_, H_2_, and CO_2_) emission per se (g/d) data were collected continuously throughout the study. The yield of CH_4_, H_2_, and CO_2_ was calculated using the total DMI (PMR + GreenFeed bait) consumed during the respective wk and was expressed as g/ kg total DMI. Similarly, methane intensity was calculated on a BW basis as g CH_4_/ kg BW.

### Feed Samples and Chemical Analysis

During the study, maize silage samples were collected daily and dry ration component samples were collected weekly to determine DM content. This information was subsequently used to determine how much of each ration component as fresh product was required to prepare the diets. Furthermore, samples of the separate PMR components (i.e., maize silage, lucerne, concentrates, and soybean meal) and GreenFeed bait, were collected weekly and stored at −20 °C pending analysis. These feed samples were sent to the laboratory of Animal Nutrition (Wageningen University & Research, Wageningen, the Netherlands) for analyses. Additionally, a representative sample of PMR was also collected from each treatment group at four different timepoints in week 3, 6, 9 and 12 of the treatment periods, and stored at −20 °C pending analysis. The PMR samples were sent to Global R&D Analytics NIC-RD/A of dsm-firmenich (Kaiseraugst, Switzerland) for the analysis of 3-NOP content, according to the procedure described by van Gastelen et al. (2020).

All collected feed samples were thawed at room temperature, freeze-dried until constant weight (i.e., maize silage only), and ground to pass a 1-mm screen by using a cross-beater mill for maize silage (Peppink 100AN) and an ultra-centrifugal mill for all other ration components (Retsch ZM200, Retsch GmbH). The samples were subsequently analyzed for DM, ash, nitrogen (**N**), starch, reducing sugars (i.e., all carbohydrates with reducing properties and soluble in 40% ethanol; except for maize silage), crude fat, neutral detergent fiber (**NDF**), acid detergent fiber (**ADF**), and acid detergent lignin (**ADL**) as described by Abrahamse et al. (2008). Bomb calorimetry (ISO 9831; International Organization for Standardization, 1998) was used to determine gross energy. Crude protein was calculated as N × 6.25, where N was determined using the Kjeldahl method with cupric sulfate as catalyst (ISO 5983; International Organization for Standardization, 2005). The N concentrations in the maize silage was determined in fresh material according to Klop et al. (2016).

### Statistical Analysis

Overall, 1 male calf needed to be physically removed from the study after week 6 of the treatment period due to aggressive behavior and data from 1 male calf was excluded after week 6 of the treatment period due to health issues (coccidiosis) compromising his DMI and BW gain. Both animals were from the 3-NOP treatment group, however, their removal from the study was not considered to be related to 3-NOP treatment. Neither of these animals could be replaced. Their data up to the point of leaving the study was included within the statistical analysis.

Statistical analysis was performed by using SAS version 9.4 (SAS Institute Inc., Cary, NC). Data was averaged for the covariate period and for treatment wk 1-3, 4-6, 7-9 and 10-12. The covariate period was used to calculate a covariate mean for all variables, which was added in the statistical models as baseline measurement. A repeated measurements model (PROC MIXED) with animal as the repeated subject was used to test the effects of treatment (CTL or 3-NOP), period (wk 1-3, 4-6, 7-9, and 10-12), their interaction, and the covariate mean on dependent variables related to emissions, visits to the GreenFeed system, feed intake, and BW. Moreover, a linear mixed model (PROC MIXED) was used to test the effects of treatment and, when applicable, the covariate mean on the dependent variables initial BW, final BW, BW gain, ADG, feed to gain ratio, and feed efficiency. For initial BW, feed to gain ratio, and feed efficiency, no covariate mean was available. For final BW, BW gain, and ADG the initial BW was used as the covariate mean. All analyses included a random effect of block and animal.

Model residuals were assessed for normality, outliers, and independence. For the number of GreenFeed visits, residuals followed a non-normal distribution. Consequently, this variable was inverse transformed for the statistical analysis, and *P*-values were obtained via bootstrapping. For the repeated measurement models, the covariance structure with the best fit was selected based on the model with the lowest corrected Akaike Information Criterion with a correction for small sample sizes (AICC). Covariance structures considered were Compound Symmetry (CS), Heterogeneous CS (CSH), Unstructured (UN), First Order Autoregressive (AR(1)), Heterogeneous AR(1) (ARH(1)). Values are presented as covariate-adjusted least squares means (**LSM**) ± pooled SEM. Significance of effects was declared at P ≤ 0.05 and trends at 0.05 < P ≤ 0.10.

## Results

### Dosage of 3-NOP

The 3-NOP dose in the PMR samples of the CTL diet was 0 mg/kg. The 3-NOP dose in the PMR samples of the 3-NOP diet was 145 mg/ kg DM, which was below target (i.e., 175 mg/ kg DM). Considering the intake of GreenFeed bait, the average level of 3-NOP in the total daily DMI was 121 mg 3-NOP/ kg DM on complete ration level, as opposed to the target dose of 150 mg 3-NOP / kg DM.

### Effect of 3-NOP on Gas Emission

For the visits to the GreenFeed system, a treatment × period interaction (P = 0.049) was observed (Table 3). In every period, calves receiving 3-NOP visited the GreenFeed system more often compared to calves receiving CTL. Measures from the GreenFeed system covered the full 24h period (Figure 1A).

**Figure 1.**
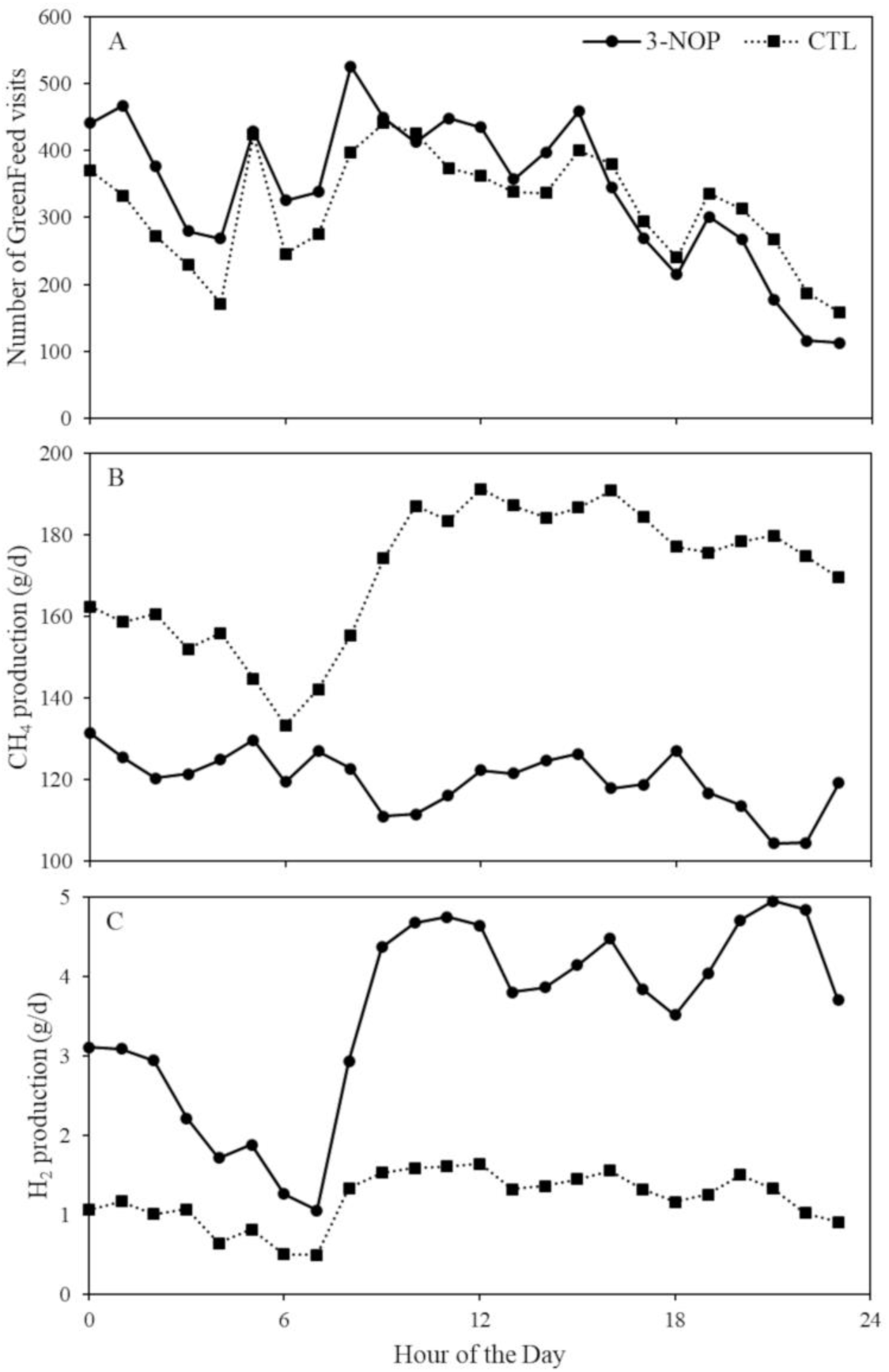
Total number of GreenFeed visits over the complete treatment period per hour (A), average CH_4_ production (g/d) per hour (B), and average H_2_ production per hour (C) of calves from 5 mo of age receiving a diet supplemented with a placebo (CTL) or with 3-nitrooxypropanol (3-NOP).

**Table 3.**
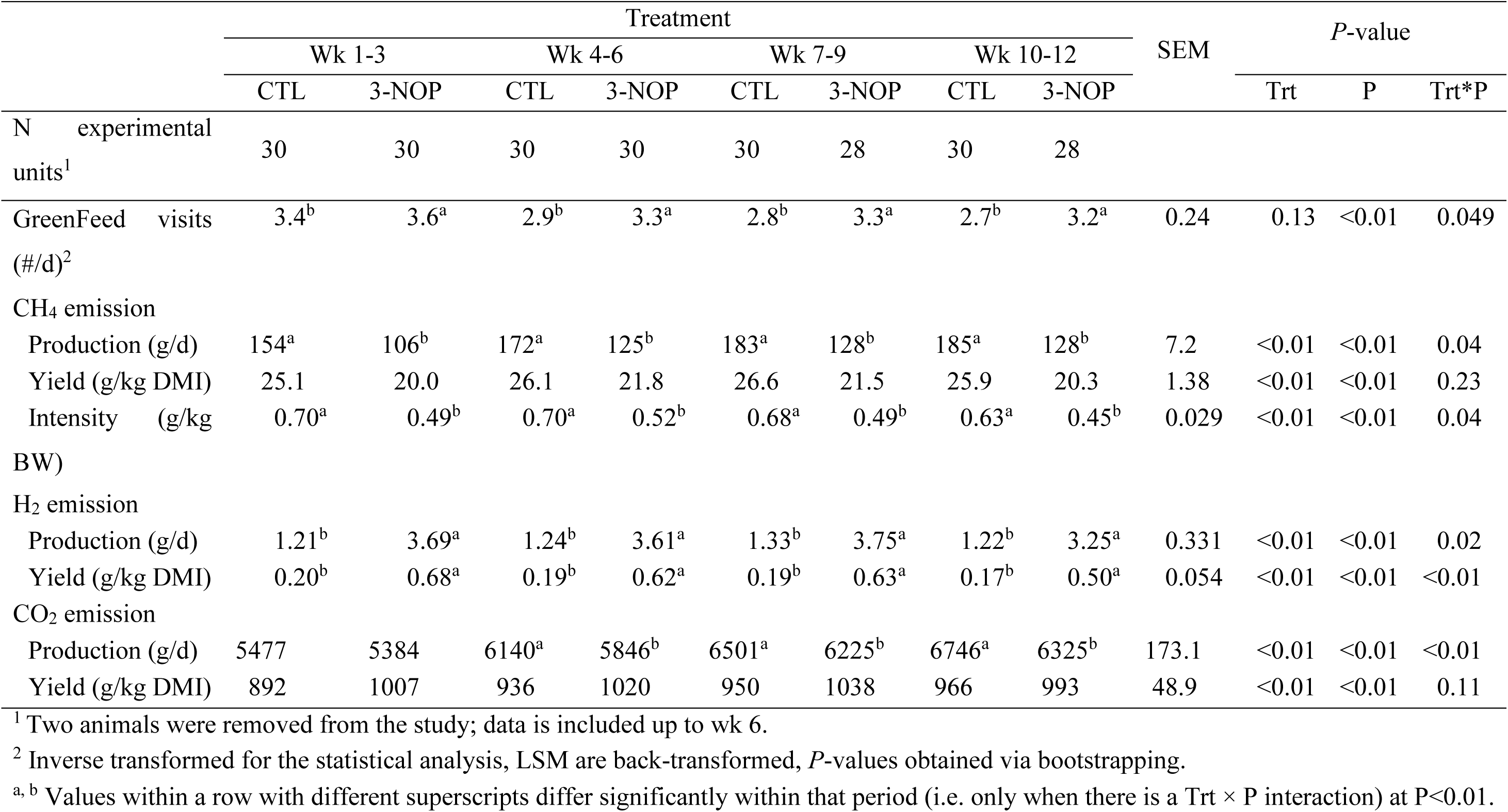
Gas emissions of growing calves from 5 mo of age receiving a diet supplemented with a placebo (CTL) or with 3-nitrooxypropanol (3-NOP) treatment (trt) per period (P; wk 1-3, wk 4-6, wk 7-9, wk 10-12) (LSM ± pooled SEM)

For the production and intensity of CH_4_, a treatment × period interaction (P = 0.04) was observed. In every period, calves receiving 3-NOP had a lower CH_4_ production and intensity compared to calves receiving CTL. Compared to CTL, CH_4_ production was reduced by 31.3% in wk 1 – 3, by 27.1% in wk 4 – 6, by 30.0% in wk 7 – 9, and by 30.7% in wk 10 – 12 for 3-NOP. Compared to CTL, CH_4_ intensity was reduced by 30.0% in week 1 – 3, by 25.8% in wk 4 – 6, by 27.9% in wk 7 – 9, and by 28.6% in wk 10 – 12 for 3-NOP. Overall, the 3-NOP calves had a reduced CH_4_ production and intensity (CTL vs. 3-NOP; 174 ± 2.0 g/d vs. 122 ± 2.0 g/d, P<0.01 and 0.68 ± 0.008 g/kg BW vs. 0.49 ± 0.008 g/kg BW, P<0.01) compared to CTL. As such, the 3-NOP calves had an overall reduction of 29.8% for CH_4_ production and 27.9% for CH_4_ intensity. No interaction between treatment and period was observed for CH_4_ yield. Overall, calves receiving 3-NOP had a reduced CH_4_ yield (CTL vs. 3-NOP; 25.9 ± 0.38 g/ kg total DMI vs. 20.9 ± 0.38 g/ kg total DMI, P<0.01), indicating a 19.4% reduction. When investigating the CH_4_ production over a 24h period, fluctuations existed, where especially during the day CH_4_ production was higher, mainly for the CTL group (Figure 1B).

For the production and yield of H_2_, a treatment × period interaction (P=0.02 and P<0.01, respectively) was observed. In every period, calves receiving 3-NOP had a greater H_2_ production and yield compared to calves receiving CTL. Compared to CTL, H_2_ production was increased with 305% in wk 1 – 3, with 291% in wk 4 – 6, with 282% in wk 7 – 9, and with 266% in wk 10 – 12 for 3-NOP. Compared to CTL, H_2_ yield was increased with 340% in wk 1 – 3, with 326% in wk 4 – 6, with 332% in wk 7 – 9, and with 294% in wk 10 – 12 for 3-NOP. Over the complete treatment period, the 3-NOP calves had an increased H_2_ production and yield (CTL vs. 3-NOP; 1.25 ± 0.095 g/d vs. 3.57 ± 0.095 g/d, P<0.01 and 0.19 ± 0.015 g/kg DMI vs. 0.61 ± 0.015 g/kg DMI, P<0.01) compared to CTL. As such, the 3-NOP calves had an overall increase of 285% for H_2_ production and 321% for H_2_ yield. When investigating the H_2_ production over a 24h period, fluctuations existed, where especially during the day H_2_ production was higher, mainly for the 3-NOP group (Figure 1C).

For the production of CO_2_, a treatment × period interaction (P<0.01) was observed. The production of CO_2_ did not significantly differ between CTL and 3-NOP in the first period (week 1-3) but was higher for CTL compared to 3-NOP in the three following periods (i.e., week 4-6, 7-9, and 10-12). Overall, calves receiving 3-NOP had a reduced CO_2_ production of 4.4% (CTL vs. 3-NOP; 6216 ± 45.3 g/d vs. 5945 ± 45.7 g/d, P<0.01), but an increased CO_2_ yield of 8.4 (CTL vs. 3-NOP; 936 ± 10.6 / kg total DMI vs. 1014 ± 10.8 g/ kg total DMI, P<0.01) during the complete treatment period.

### Effect of 3-NOP on Feed Intake and Body Weight

For DMI of GreenFeed bait, a treatment × period interaction (P<0.01) was observed, where during all periods, calves receiving 3-NOP had a greater DMI of GreenFeed bait compared to calves receiving CTL (Table 4) (P<0.01 for all comparisons between the groups within the same period). During the complete treatment period, calves receiving 3-NOP had a lower DMI of PMR (5.0 ± 0.07 kg/d) compared to calves receiving CTL (5.9 ± 0.07 kg/d, P<0.01). Moreover, overall, the calves in the 3-NOP group had a lower total DMI compared to the calves in the CTL group (6.1 ± 0.08 kg/d vs. 6.9 ± 0.07 kg/d, P<0.01).

**Table 4.**
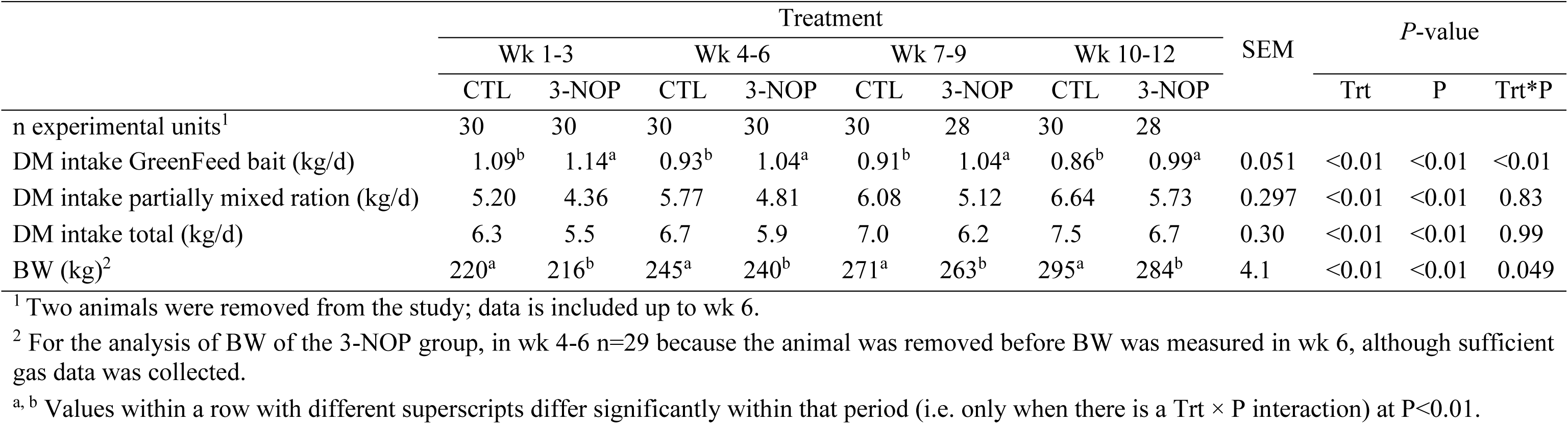
Dry matter intake and BW of growing calves from 5 mo of age receiving a diet supplemented with a placebo (CTL) or with 3-nitrooxypropanol (3-NOP) treatment (trt) per period (P; wk 1-3, wk 4-6, wk 7-9, wk 10-12) (LSM ± pooled SEM)

|  | Treatment |  |  |  |  |  |  |  | SEM | P-value |  |  |
| --- | --- | --- | --- | --- | --- | --- | --- | --- | --- | --- | --- | --- |
|  | Wk 1-3 |  | Wk 4-6 |  | Wk 7-9 |  | Wk 10-12 |  |  | Trt | P | Trt*P |
|  | CTL | 3-NOP | CTL | 3-NOP | CTL | 3-NOP | CTL | 3-NOP |  |  |  |  |
| n experimental units<sup>1</sup> | 30 | 30 | 30 | 30 | 30 | 28 | 30 | 28 |  |  |  |  |
| DM intake GreenFeed bait (kg/d) | 1.09<sup>b</sup> | 1.14<sup>a</sup> | 0.93<sup>b</sup> | 1.04<sup>a</sup> | 0.91<sup>b</sup> | 1.04<sup>a</sup> | 0.86<sup>b</sup> | 0.99<sup>a</sup> | 0.051 | <0.01 | <0.01 | <0.01 |
| DM intake partially mixed ration (kg/d) | 5.20 | 4.36 | 5.77 | 4.81 | 6.08 | 5.12 | 6.64 | 5.73 | 0.297 | <0.01 | <0.01 | 0.83 |
| DM intake total (kg/d) | 6.3 | 5.5 | 6.7 | 5.9 | 7.0 | 6.2 | 7.5 | 6.7 | 0.30 | <0.01 | <0.01 | 0.99 |
| BW (kg)<sup>2</sup> | 220<sup>a</sup> | 216<sup>b</sup> | 245<sup>a</sup> | 240<sup>b</sup> | 271<sup>a</sup> | 263<sup>b</sup> | 295<sup>a</sup> | 284<sup>b</sup> | 4.1 | <0.01 | <0.01 | 0.049 |

<sup>1</sup> Two animals were removed from the study; data is included up to wk 6.
<sup>2</sup> For the analysis of BW of the 3-NOP group, in wk 4-6 n=29 because the animal was removed before BW was measured in wk 6, although sufficient gas data was collected.
<sup>a, b</sup> Values within a row with different superscripts differ significantly within that period (i.e. only when there is a Trt × P interaction) at P<0.01.

For BW, a treatment × period interaction (P=0.049) was observed, also indicating that calves in the 3-NOP group gained less BW during the treatment period than the calves of the CTL group. Overall, calves in the 3-NOP group had a lower BW compared to CTL (251 ± 1.2 kg vs 258 ± 1.2 kg, P<0.01). The 3-NOP group and the CTL group did not differ in initial BW, but calves in the 3-NOP group had a lower final BW compared to CTL (Table 5). As a result, calves in the 3-NOP group also had a lower BW gain and lower ADG during the complete treatment period compared to CTL. Despite the observed difference in DMI and ADG between the 3-NOP and CTL group, feed efficiency was not significantly affected by 3-NOP supplementation.

**Table 5.** Body weight and feed efficiency of growing calves from 5 mo of age receiving a diet supplemented with a placebo (CTL) or with 3-nitrooxypropanol (3-NOP) (LSM ± pooled SEM)

|  | Treatment |  | SEM | *P*-value |
| --- | --- | --- | --- | --- |
|  | CTL | 3-NOP |  | Trt |
| n experimental units<sup>1</sup> | 30 | 28 |  |  |
| BW initial (kg)<sup>2</sup> | 203.1 | 202.4 | 4.96 | 0.83 |
| BW final (kg) | 300.5<sup>a</sup> | 290.3<sup>b</sup> | 3.22 | <0.01 |
| BW gain (kg) | 98.2<sup>a</sup> | 87.9<sup>b</sup> | 3.22 | <0.01 |
| Average daily gain (kg) | 1.17<sup>a</sup> | 1.05<sup>b</sup> | 0.04 | <0.01 |
| Feed to gain ratio | 5.90 | 5.84 | 0.236 | 0.77 |
| Feed efficiency<sup>3</sup> | 0.173 | 0.175 | 0.0061 | 0.61 |

<sup>1</sup> Two animals in the 3-NOP group were removed from the study.
<sup>2</sup> For the analysis of the initial BW of the 3-NOP group, n=29: excluded the datapoint of only one animal, because data was excluded from this animal in the latter part of the study due to health issues after wk 6. For the other variables in this table, both animals that were removed after wk 6 are excluded (i.e., n=28).
<sup>3</sup> Gain to feed ratio.
<sup>a, b</sup> Values within a row with different superscripts differ significantly at $P < 0.05$ .

## Discussion

### Methane Emission

This study aimed to assess the efficacy and safety of 3-NOP for growing calves from 5 mo of age. Considering the efficacy, we hypothesized that 3-NOP supplementation would decrease CH_4_ emission and that this effect would be persistent over time. Indeed, in the current study, supplementation with 3-NOP reduced CH_4_ production, yield, and intensity by 29.8%, 19.4%, and 27.9% compared to CTL, respectively, with relatively minor differences between periods (< 5% units). These reduction percentages were slightly lower than what could be expected based on the meta-analysis by Orzuna-Orzuna et al. (2024; production: 34.9% and yield: 29.6%), but slightly higher than what could be expected based on the meta-analysis by Dijkstra et al. (2018; production: 22.2% and yield: 17.1%, after adjustment for the effects of 3-NOP dose and dietary NDF content). Overall, CH_4_ production was 174 g/d for calves receiving CTL and 122 g/d for calves receiving 3-NOP, which was lower compared to Alemu et al. (2021) on growing beef steers with an average start BW of 282 kg, and Vyas et al. (2018) on beef cattle with an average start BW of 308 kg. The calves in the current study were younger than the animals used in the above-mentioned studies and, likely related, had a lower BW and DMI, explaining the lower CH_4_ production values. Also, differences in dietary composition explains a part of the difference in CH_4_ production levels as well.

Despite the persistent CH_4_ reduction by 3-NOP, an interaction effect between treatment and period was observed for both CH_4_ production and the intensity. However, within each period, both CH_4_ production (−31.3% for wk 1-3, −27.1% for wk 4-6, −30.0% for wk 7-9, and −30.7% for wk 10-12) and intensity (−30.0% for wk 1-3, −25.8% for wk 4-6, −27.8% for wk 7-9, and −28.6% for wk 10-12) were lower for calves receiving 3-NOP compared to CTL, suggesting a persistent CH_4_ reducing effect of 3-NOP. The animals enrolled were growing calves from 5 mo of age, which implies that the DMI and BW of the calves increased over the course of the study, and as expected, CH_4_ production also increased. Especially for the CTL calves, CH_4_ production increased during the treatment period, explaining the observed interaction between treatment and period for CH_4_ production. Due to a similar increase in CH_4_ production and DMI over time, CH_4_ yield was stable over time for both CTL and 3-NOP calves, explaining the lack of interaction for CH_4_ yield. In contrast, as the increase in BW over time was slightly greater compared to the increase in CH_4_ production, the CH_4_ intensity declined only slightly over time. The greater increase in BW for calves receiving CTL compared to 3-NOP may have resulted in the interaction between treatment and period. The meta-analysis by Orzuna-Orzuna et al. (2024) demonstrated that the daily CH_4_ emission decreased for calves receiving 3-NOP, regardless of the supplementation period. In line with these results, in the current study, the reductions of CH_4_ production, yield, and intensity did not decrease over the course of the treatment duration, indicating a persistent efficacy of 3-NOP on the reduction of CH_4_ emission, and no adaptation to 3-NOP supplementation over the (relatively short) treatment period of 12 wk.

### Hydrogen and Carbon Dioxide Emission

Overall, the H_2_ production was 3-fold greater and H_2_ yield was over 3-fold greater in the calves receiving 3-NOP compared to CTL. This increase in H_2_ is often observed when methanogenesis – the main H_2_ sink – is inhibited via 3-NOP supplementation (e.g., Vyas et al., 2016; Alume et al., 2021). However, the percentage increase in both H_2_ production and yield (with the latter only a slight change) reduced over the course of the treatment period. This reduction in H_2_ emission was unrelated to the percentage reduction for both CH_4_ production and yield upon 3-NOP supplementation. This suggests that with time of 3-NOP application, H_2_ may have been redirected into an alternative H_2_ sink, because H_2_-consuming pathways are often upregulated when ruminal H_2_ concentration is increasing (Jansen, 2010), and to propionate and butyrate at the expense of acetate delivering less H_2_. This increase in propionate and butyrate upon 3-NOP supplementation has been observed in other studies with maize silage-based diets (e.g., Romero-Perez et al., 2014; Haisan et al., 2016). The measured increase in H_2_ emission upon 3-NOP supplementation was lower than the expected stoichiometric amount that involved with the decrease in CH_4_ production. This lower-than-expected recovery of H_2_ that was spared from utilization with methanogenesis is also observed in other studies, including Hristov et al. (2015) and van Gastelen et al. (2020; 2022). Partly, it may have been caused by the measurement technique that was used in our study. By using the GreenFeed system as a spot sampling device, we may have missed the rapidly occurring peaks in H_2_ emission. Van Lingen et al. (2017) showed that H_2_ emissions were a factor 100 greater in the first hour after a meal. In line with this postprandial response of H_2_ emission, de Mol et al. (2024) demonstrated that the level of H_2_ measured by the GreenFeed system was affected by the time interval between a preceding meal and the actual H_2_ measurement (i.e. level of H_2_ production was higher with a shorter time interval). In the present study, fresh feed was usually presented between 0800 and 0900 h and the highest frequency of visits to the GreenFeed also occurred around that time and directly thereafter. However, some calves might have consumed larger quantities of feed later during the day as not all animals could access the fresh feed at the same time (i.e., 6 FB for 30 calves). If the calves did not visit the GreenFeed system shortly after consuming a meal at the FB, the large peak in H_2_ emission following a meal is not accounted for in the daily estimate. Hence, H_2_ emission could have been underestimated. Van Lingen et al. (2023) demonstrated that a sampling interval of 2.0h was required to obtain accurate daily H_2_ emission values of dairy cows fed twice daily ad libitum to make them not differ from continuous measurements in climate respiration chambers. Although GreenFeed measurements covered a full 24h period in the current study (Figure 1A), the calves were only allowed to visit the GreenFeed system four times per day, leading to an interval period of 8h which is considerably larger than the 2.0h interval as suggested by van Lingen et al. (2023). In correspondence with this, much higher H_2_ emissions are measured in climate respiration chambers than with GreenFeed systems.

In the current study, calves receiving 3-NOP had a lower CO_2_ production compared to the CTL calves from wk 4 onwards. However, overall, calves receiving 3-NOP had an increased CO_2_ yield compared to CTL. This was probably due to a lower DMI in the 3-NOP supplemented animals, effectively leading to a higher CO_2_ yield but also to a slower growth rate, and to a small extent perhaps by a lower CO_2_ utilization due to less CH_4_ production. In the current study, overall CO_2_ production was lower (6.1 kg/d) compared to the meta-analysis by Orzuna-Orzuna et al. (2024; 8.5 kg/d), likely because the calves in our study had a younger age, and subsequently a lower BW and DMI than those included in the meta-analysis. The yield of CO_2_ was however slightly higher (975 g/kg DMI) compared to the meta-analysis of Orzuna-Orzuna et al. (2024; 865 g/kg DMI). The relative difference between our study and the meta-analysis was greater for DMI than for CO_2_ production, resulting in the somewhat higher CO_2_ yield of the calves in our study.

### Feed Intake and Body Measures

Supplementation with 3-NOP resulted in an overall reduction of DMI, although calves receiving 3-NOP visited the GreenFeed more often in every period and had a slightly greater GreenFeed bait intake compared to CTL calves. Hence, the reduction in DMI of calves receiving 3-NOP was primarily driven by a reduction in PMR intake. Moreover, the reduction in DMI started as soon as the treatment started for the calves receiving 3-NOP. Some other studies on growing calves also reported a reduction in DMI upon supplementation with 3-NOP (Romero-Perez et al., 2014; Vyas et al., 2016). In these 2 studies, as well as in the current study, starch-rich diets (i.e., >250 g/ kg DM) were fed. Kirwan et al. (2014) did not observe an effect on DMI upon 3-NOP supplementation in growing beef cattle, but fed a diet with a lower starch content (i.e., 189 g/ kg DM). In an exploratory analysis, potential interactions between the diet fed and the effect of 3-NOP on DMI within the current study and the study by Kirwan et al. (2024) were investigated. These analyses did not indicate a clear interaction, and possible dietary consequences for the effect of 3-NOP on DMI will need to be investigated further. When fermentable carbohydrates, such as starch, are metabolized in the rumen, volatile fatty acids (**VFA**) are produced, including propionate. According to the meta-analysis by Orzuna-Orzuna et al. (2024), ruminal concentration of propionate increases upon 3-NOP supplementation. An increased production of VFA as well as an increased propionate molar proportion could result in an increased portal vein propionate concentration, which can promote satiety, resulting in a decreased DMI (Allen, 2000; Stocks and Allen, 2012). However, the decrease in DMI could also indicate that the rumen microbiota of the calves receiving 3-NOP were not able to cope with the excess of the H_2_ that was formed because of inhibiting methanogenesis, as an increased ruminal partial pressure of H_2_ can be expected to inhibit the metabolism of rumen microorganisms (McAllister and Newbold, 2008; Morgavi et al., 2010). Although such an inhibitory effect of H_2_ cannot be excluded, it is worth reminding that normal postprandial fluctuations in rumen H_2_ are much larger (van Lingen et al., 2017).

The reduced DMI for calves receiving 3-NOP resulted in a lower overall BW gain (i.e., difference initial and final BW) as well as a lower BW during all periods compared to the calves receiving CTL. In earlier studies, BW of calves receiving 3-NOP remained unaffected (Kirwan et al., 2024), even when DMI was reduced upon 3-NOP supplementation (Romero-Perez et al., 2014; Vyas et al., 2016; Orzuna-Orzuna et al., 2024). In the current study, BW was directly affected by the drop in DMI, and this was consistent throughout the entire measurement period. A lower DMI means that there is less energy available for growth. Moreover, a lower DMI results in less feed in the gastrointestinal tract, per definition resulting in a lower BW. The lower BW in this study suggests that the potential benefit from decreased CH_4_ emission (i.e., less loss of energy) did not outbalance the negative impact of the decrease in DMI, whereas this may have been the case in other studies where growth performance was equal with 3-NOP treatment. The increased proportion of ruminal propionate upon feeding 3-NOP (Orzuna-Orzuna et al., 2024) may yet again be part of the explanation. As propionate is a glucogenic nutrient, it may result in increased insulin blood levels reducing the mobilization of body reserves (van Knegsel et al., 2007). As a result, fat deposition could be enhanced rather than muscle accumulation, leading to a lower BW gain per unit of energy retained due to water associated with muscle protein. In the current study, supplementation with 3-NOP did not significantly affect feed efficiency or feed to gain ratio, indicating that any reduction in BW was a direct result of a reduction in DMI.

## Conclusion

Feeding 121 mg 3-NOP/ kg DM on complete ration level to growing calves over a 12 wk period significantly and persistently reduced the enteric CH_4_ production (g/d) by 29.8%, yield (g/k DMI) by 19.4%, and intensity (g/kg BW) by 27.8%. This was accompanied with a (less persistent) 3-fold increase in H_2_ emission, and lower BW gain (−10.3 kg over 12-wk trial period) and total DMI (−0.8 kg/d), but unaffected feed efficiency. It can be concluded that 3-NOP is a promising strategy to persistently decrease CH_4_ emission in growing beef calves from 5 to 8 mo of age, without negatively impacting feed efficiency.

## Abbreviations

ADG: average daily gain
ADF: acid detergent fiber
ADL: acid detergent lignin
BW: body weight
CH_4_: methane
CO_2_: carbon dioxide
CTL: control diet with placebo concentrate
DM: dry matter
DMI: dry matter intake
FB: feed bin
GHG: greenhouse gas
H_2_: hydrogen
N: nitrogen
NDF: neutral detergent fiber
PMR: partial mixed ration
LSM: least squares means
VFA: volatile fatty acid
3-NOP: 3-nitrooxypropanol.

## Acknowledgements

The staff of Dairy Campus, in particular Martin de Bree, Paul Huisman, Fedde de Jong, and Anna Klaver are acknowledged for their assistance during the implementation of the experiment. Furthermore, the laboratory staff of the Animal Nutrition Group (Wageningen, the Netherlands) is acknowledged for all chemical analyses. The study was financed by dsm-firmenich (Basel, Switzerland).

